# Catestatin selects for the colonization of antimicrobial-resistant gut bacterial communities

**DOI:** 10.1101/2021.10.11.463921

**Authors:** Pamela González-Dávila, Markus Schwalbe, Arpit Danewalia, Boushra Dalile, Kristin Verbeke, Sushil K Mahata, Sahar El Aidy

## Abstract

The gut microbiota is in continuous interaction with the innermost layer of the gut, namely the epithelium. One of the various functions of the gut epithelium, is to keep the microbes at bay to avoid overstimulation of the underlying mucosa immune cells. To do so, the gut epithelia secrete a variety of antimicrobial peptides, such as chromogranin A (CgA) peptide catestatin (CST: hCgA_352-372_). As a defense mechanism, gut microbes have evolved antimicrobial resistance mechanisms to counteract the killing effect of the secreted peptides. To this end, we treated wild-type mice and CST knockout (CST-KO) mice (where only the 63 nucleotides encoding CST have been deleted) with CST for 15 consecutive days. CST treatment was associated with a shift in the diversity and composition of the microbiota in the CST-KO mice. This effect was less prominent in WT mice. Levels of the microbiota-produced short-chain fatty acids, in particular, butyrate and acetate were significantly increased in CST-treated CST-KO mice but not the WT group. Both CST-treated CST-KO and WT mice showed a significant increase in microbiota-harboring phosphoethanolamine transferase-encoding genes, which facilitate their antimicrobial resistance. Finally, we show that CST was degraded by *Escherichia coli* via an omptin-protease and that the abundance of this gene was significantly higher in metagenomic datasets collected from patients with Crohn’s disease but not with ulcerative colitis. Overall, this study illustrates how the endogenous antimicrobial peptide, CST, shapes the microbiota composition in the gut and primes further research to uncover the role of bacterial resistance to CST in disease states such as inflammatory bowel disease.

## Introduction

The complex microbial community inhabiting the mammalian gut, and in particular their metabolic products, has a crucial effect on various health and disease states, such as inflammatory bowel disorders ^1^. Gut microbiota, and its products, are in continuous interaction with the innermost layer of the gut, namely the gut epithelium. Part of the gut epithelial function is to keep the microbes at bay to avoid overstimulation of the underlying immune cells in the gut, and the subsequent inflammation ^2,3^. To do so, the gut epithelial cells secrete a wide variety of antimicrobial peptides.

CST, a proteolytically processed product of Chromogranin-A (CgA), acts as an antimicrobial peptide ^4,5^. The pro-protein CgA is abundantly expressed in endocrine and neuroendocrine cells including the epithelial enteroendocrine cells (EECs) ^6–9^. CST consists of 21 amino acid residues and acts on a wide range of regulatory functions including immune, endocrine, metabolic, neurological, and cardiovascular functions ^10–14^. For example, mice with a selective deletion of the CST-coding region of the *Chga* gene (CST-KO mice) displayed increased gut permeability, which was restored by CST treatment ^15^.

To date, a few studies have indicated associations between CST and the gut microbiota. Intrarectal administration of CST for 6 days, in mice, resulted in a shift in the gut microbiota composition, with a decrease of *Firmicutes* and an increase in the *Bacteroidota* phyla (Rabbi, Munyaka, and Eissa 2017). These findings suggest the role of CST as a modulator of the gut microbiota composition. This CST-modulatory effect could be mediated by its antimicrobial activity against a wide range of microorganisms. In fact, CST has been reported to have antimicrobial activity against pathogenic bacteria and filamentous fungi in *in-vitro* studies, ^5^. In humans, CST showed antimicrobial effects against several skin microbes ^16^. Nevertheless, the underlying mechanisms by which CST exerts its effects *in vivo* on the gut microbiota remain obscure. In this study, we show how CST plays a role in governing the colonization of the gut microbiota.

## Results

### Catestatin treatment affects the microbiota composition in CST-KO and WT mice

The composition of the gut microbiota is shaped by, among others, host-produced antimicrobial peptides ^17,18^. Here, we treated CST-KO and WT mice (n = 12) with 2 μg/g body weight/day CST by intraperitoneal injection for 15 consecutive days. Amplicon sequencing of the V3-V4 regions of the bacterial 16S rRNA gene was performed on the fecal samples of CST-KO and WT mice with and without CST treatment (n = 12). Microbial richness, assessed by the number of observed amplicon sequence variants (ASVs) and ACE index (Abundance-based Coverage Estimator), showed a significant decrease in CST-KO compared to WT mice **(Figure 1A)**. In contrast, the richness levels were restored upon treatment of CST-KO mice, but not in WT mice treated with CST. Next, the microbiota diversity was determined by Shannon’s *H* and inverted Simpson’s index, both indices are used to measure similar parameters of alpha diversity. Similar to the richness scores, the diversity index was significantly higher in the CST-treated CST-KO mice but not in the CST-treated WT group **(Figure 1B)**. The data highlight that the effect of the absence of CST in CST-KO mice on the microbiota diversity and richness could be restored upon treatment with CST, while in WT mice, with normal CST levels in their gut, the microbiota diversity and richness did not change with CST treatment.

**Figure 1.**
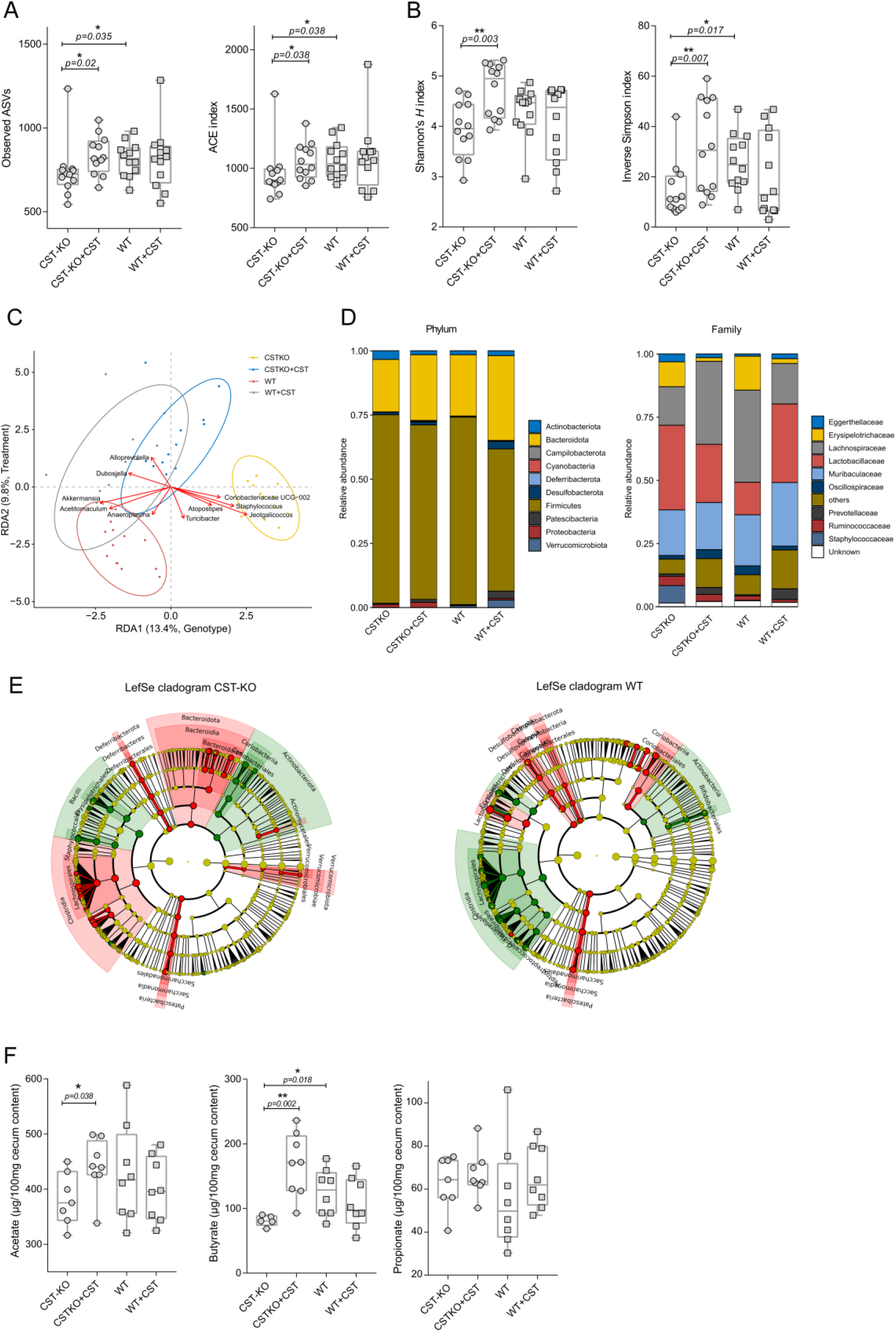
CST treatment affects the murine fecal microbiota. (**A, B**) Alpha diversity was assessed using different metrics and is concordantly significantly increased upon CST treatment. **(A)** represent the species richness (observed ASVs, ACE) and (**B**) species diversity of the CST-KO and WT mice before and after CST treatment. Significance was tested with an unpaired *Mann-Whitney* test. Boxes represent the median with interquartile range, and whiskers represent the maxima and minima. (**C**) Genotype and treatment constrained redundancy analysis (RDA) on genus collapsed abundances. Arrows indicate the association of taxa with samples, with the length being a proxy for the strength of the association. Ellipses represent normal data generated by the R package ggplot2. Significant separation of clusters and contribution of the variables to the variance of the RDA was tested with Permutational Multivariate ANOVA (PERMANOVA) and revealed significant (p < 0.001) effects of CST administration, both in CST-KO and WT mice; grey – WT+CST, yellow – CST-KO, blue - CST-KO+CST. (**D**) Relative abundance of the present phyla and families. (**E**) Cladograms of a LefSe analysis (Linear discriminant analysis Effect Size) comparing microbiota changes upon CST treatment in CST-KO and WT mice. Significance was tested using a Kruskal-Wallis (KW) test and Linear Discriminant Analysis (LDA). A significant feature was considered when *p* < 0.01 and Log (LDA score) > 3. The green color indicates the microbiota composition before CST treatment and the red color after CST treatment. (**F**) Concentrations of cecal acetate, butyrate, and propionate, of CST-treated (CST-KO, n = 7; WT, n=8) and untreated animals (CST-KO, n=8; WT, n=8). Cecal acetate and butyrate were significantly increased in CST-treated CST-KO mice. Boxes represent the median with interquartile range, and whiskers represent the maxima and minima. Data were analyzed using a 2-tailed paired *t*-test, while 2 outliers were removed using ROUT in GraphPad Prism. Error bars are shown as mean ± SEM.

As a general exploratory analysis, principal component analysis (PCA) was performed and showed distinct clustering of the CST-KO, WT, and CST-treated groups **(Supplementary Figure 1A)**. Particularly, the CST-KO group clustered away from the other 3 groups, while the cluster of CST-treated WT mice showed a great spread and was overlapping with clusters of CST-treated CST-KO mice and WT mice.

To correct for any residual variation in the data, we further employed constrained RDA (redundancy analysis) to determine which bacterial taxa were associated with the different groups of mice at the genus level. The analysis was constrained to both, the mouse genotype and treatment (non-treated and CST-treated). Similar to the cluster separation detected by PCA analysis **(Supplementary Figure 1A)**, both constraints had a significant influence on the model (*p* < 0.001, determined by ANOVA-like permutation test), explaining 13.4% (genotype) and 9.8% (treatment) of the variation, respectively (**Figure 1C**). Notably, the CST treated CST-KO group was shifted to the left side along RDA1, indicating a shift in “genotype” upon CST treatment, as opposed to WT, where the treatment group only moved along RDA2, which is associated with the treatment.

To assess the treatment effect exclusively on each genotype, we performed RDA constrained to untreated and CST-treated groups. On the genus level, *Dubosiella* and *Romboutsia* showed the strongest association with CST treatment along RDA1 (explaining ~31.2% of the variation) in the CST-KO mice (**Supplementary Figure 1B**). In WT mice, RDA demonstrated an association of *Faecalibaculum, Bifidobacterium, Romboutsia*, and *Anaeroplasma* with the baseline along RDA1 (~17.2% explained variance), while *Akkermansia* was more associated with PC1 (~25.1% explained variance; residual variance). *Alloprevotella* and *Candidatus Saccharimonas* representing the strongest associations with CST treatment in WT (**Supplementary Figure 1B**). To further identify which bacterial taxa were affected by the CST treatment, pairwise comparisons of bacterial abundances were performed between CST-treated and untreated groups for each genotype. Focusing on the phylum level, Firmicutes decreased in relative abundance, while Bacteroidota, Patescibacteria, Desulfobacterota, and Proteobacteria increased with CST treatment in both CST-KO and WT groups. Verrucomicrobiota showed lower abundance in CST-KO but increased significantly in CST-treated groups **(Figure 1D)**. The changes depicted on the phyla level in CST-treated WT mice are consistent with findings from Rabbi et al, albeit different CST administration and dosage regimen ^19^. These shifts in the main bacterial phyla upon CST treatment was further confirmed to be consistent in human microbiota by *in vitro* culturing of healthy human fecal samples with 10 μM CST **(Supplementary Figure 1C)**. On family level the most prominent changes were observed for Erysipelotrichaceae, which was consistently decreased in treated groups and Lachnospiraceae, which was increased in CST-KO, but decreased in WT in the CST treated groups. Further Lactobacillaceae was found to be decreased in CST-KO, but increased in WT CST treated groups **(Figure 1D)**. On the genus level, CST treatment reduced the abundance of *Staphylococcus* and *Turicibacter* in CST-KO and WT mice, while *Alistipes, Akkermansia, and Roseburia*, were significantly increased only in the CST-KO group (**Supplementary Excel Sheet**). Next, we used LEfSe (Linear discriminant analysis Effect Size; Segata et al., 2011) to complement our differential abundance analysis. The main discriminant feature separating the groups (untreated and CST-treated) in CST-KO mice were species from the genera *Atopostipes, Jeotgalicoccus, Turicibacter, Staphylococcus*, and Coriobacteriaceae *UCG-002* family **(Figure 1E)**. In WT mice, the most discriminating genera were *Lactobacillus*, *Candidatus Sacchararimonas*, *Turicibacter*, and *Enterococcus*.

Changes in microbial composition are often accompanied by metabolic changes, in particular, the production of short-chain fatty acids (SCFAs) ^20^. Thus, levels of SCFAs; acetate, butyrate, and propionate, were measured in the cecum of untreated and CST-treated CST-KO and WT mice **(Figure 1F)**. CST treatment significantly increased the levels of acetate and butyrate in CST-KO mice but not in the WT group. Specifically, butyrate was significantly lower in the CST-KO mice compared to their WT counterparts. Both observations are consistent with the observed significant increase in the SCFA-producing bacteria, *Alistipes*, *Akkermansia*, and *Roseburia* in the CST-KO group.

Overall, the results indicated a significant impact of CST treatment on the diversity and composition of the microbiota in the CST-KO mice, where CST is absent. This effect was much less prominent in WT mice, which have normal levels of CST.

### CST treatment of CST-KO mice promotes the growth of core taxa present in WT mice

To investigate whether certain taxa exhibit similar behavior (increase or decrease in abundance) between genotypes when CST is administered, we focused on the core microbiota (common taxa between CST-treated groups), representing microbiota members, which directly (e.g., via resistance genes) or indirectly (e.g., via cross-feeding), resist the antimicrobial effect of CST. To do so, log-fold changes between CST treated and untreated groups for both WT and CST-KO mice were compared **(Figure 2)**. Almost all significant changes in core taxa (determined by unpaired Wilcoxon test) were exclusively higher in abundance in CST treated CST-KO mice (quadrants I and II), while, for WT, both significant increases and decreases in the abundance of core taxa were observed. These results coincide with the RDA analysis (cluster of CST-treated CST-KO group moving closer to WT) and indicate that the effect of CST treatment in CST-KO mice promotes the growth of core taxa associated with WT mice. Taxa present in quadrant II, which represent those common taxa between the untreated WT and treated CST-KO mice, are partially restored in their abundance in the latter group. Of note is, that *Turicibacter* was the only core taxon in both genotypes, exclusively negatively affected by CST treatment, exhibiting a significant decrease in abundance and therefore suggesting higher susceptibility to CST. To confirm this observation, we performed *in vitro* assays, to determine minimum inhibitory concentrations (MIC) for *Turicibacter sanguinis*. MIC was also performed for *Bacteroides thetaiotaomicron*, which increased in abundance upon CST treatment. The screening revealed low MIC for *T. sanguinis* (32 μM CST), while *B. thetaiotaomicron* had a higher MIC (>64 μM CST), thus confirming our *in vivo* observations. Reverse-phase HPLC analysis of spent culture supernatant after 24 hrincubation with CST revealed degradation/uptake of CST from the medium, in the case of *B. thetaiotaomicron*, but seemingly less for *T. sanguinis*, hence explaining the higher MIC value in *B. thetaiotaomicron* (**Supplementary Figure 2**). Taken together, the data imply that the changes in the gut microbiota composition associated with the CST treatment **(Figure 1)** may be caused by the antimicrobial resistance of certain taxa to CST. However, the underlying mechanisms remain obscure.

**Figure 2.**
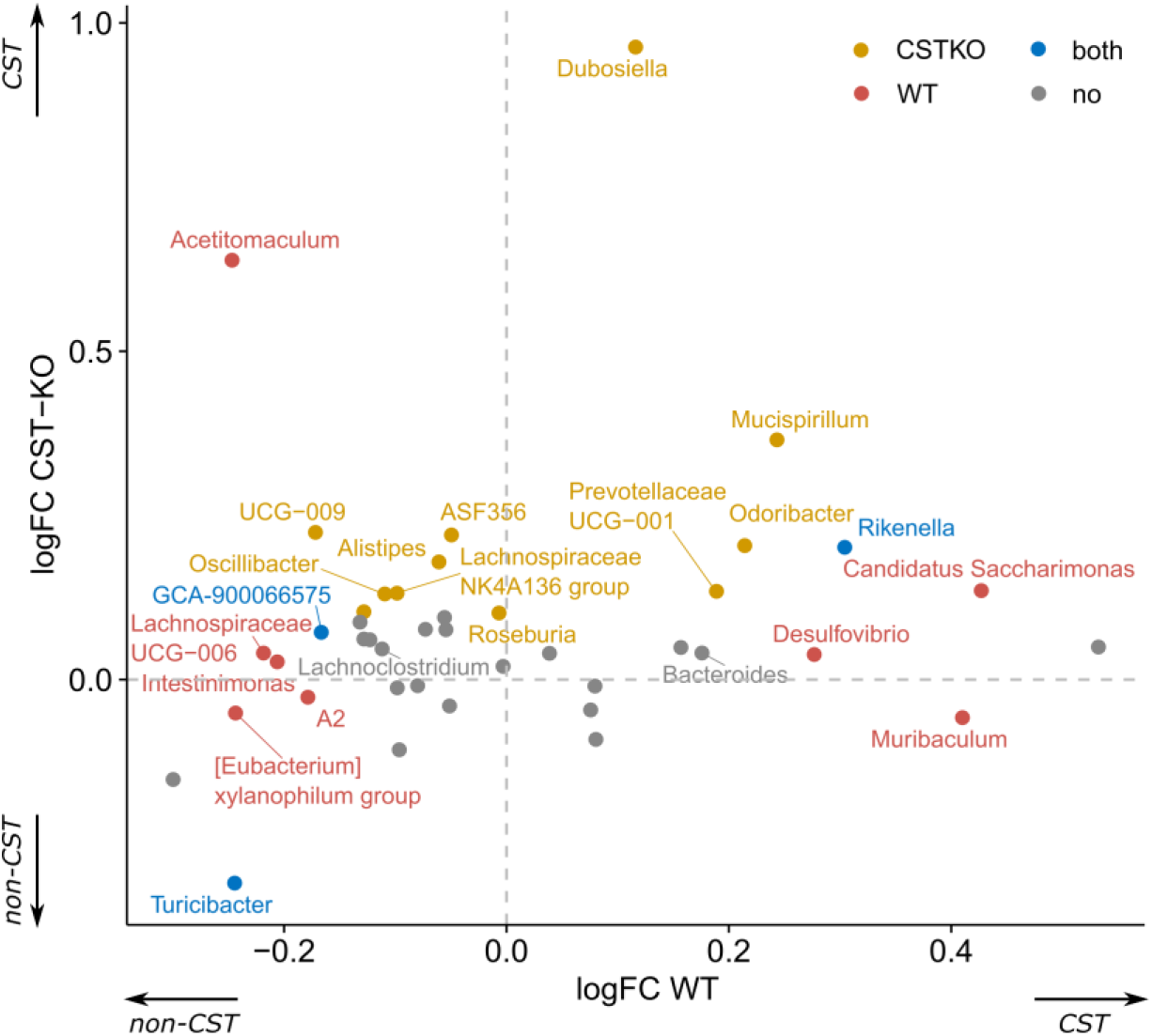
Taxa able to endure CST treatment benefit from its presence in the environment. Scatter plot of log-fold changes, comparing abundance changes for significantly abundant genera between treated and untreated groups per genotype. Colors represent significance when testing for differential abundance between treated and untreated groups for each genotype.

### Catestatin treatment promotes the abundance of the gut microbiota harboring antimicrobial resistance phosphoethanolamine transferase-encoding genes

To determine whether bacteria with an increase in abundance in CST-treated groups harbor antimicrobial resistance genes, a metagenomic prediction was performed using PiCRUSt2 on KO (KEGG Orthology) level. From a panel of known AMP resistance genes ^21^, one KEGG Orthologue (K03760) was found to be significantly different in abundance between CST treated and untreated WT mice and close to significance in CST-KO, when corrected for multiple testing (**Supplementary Excel Sheet**).

Sequences within this KEGG Orthogroup contain orthologs of EptA (*E. coli*) phosphoethanolamine transferase. As KEGG Orthologs only contain sequences from reference genomes with no specific gut metagenomic context, we sought to refine our search to investigate the taxonomical distribution of the EptA. To do so, the EptA protein sequence from *E. coli* BW25113 was used as a query to search the mouse gastrointestinal bacterial catalog (MGBC). Identified protein sequences were further filtered and a phylogenetic tree was constructed, which exhibited a distinct grouping of sequences without great dispersion and most of the sequences of the same phylum were clustered in separate clades. This showed a major subdivision between of the foremost sequences from Bacteroidota, Proteobacteria and Campylobacterota **(Figure 3A)**.

**Figure 3.**
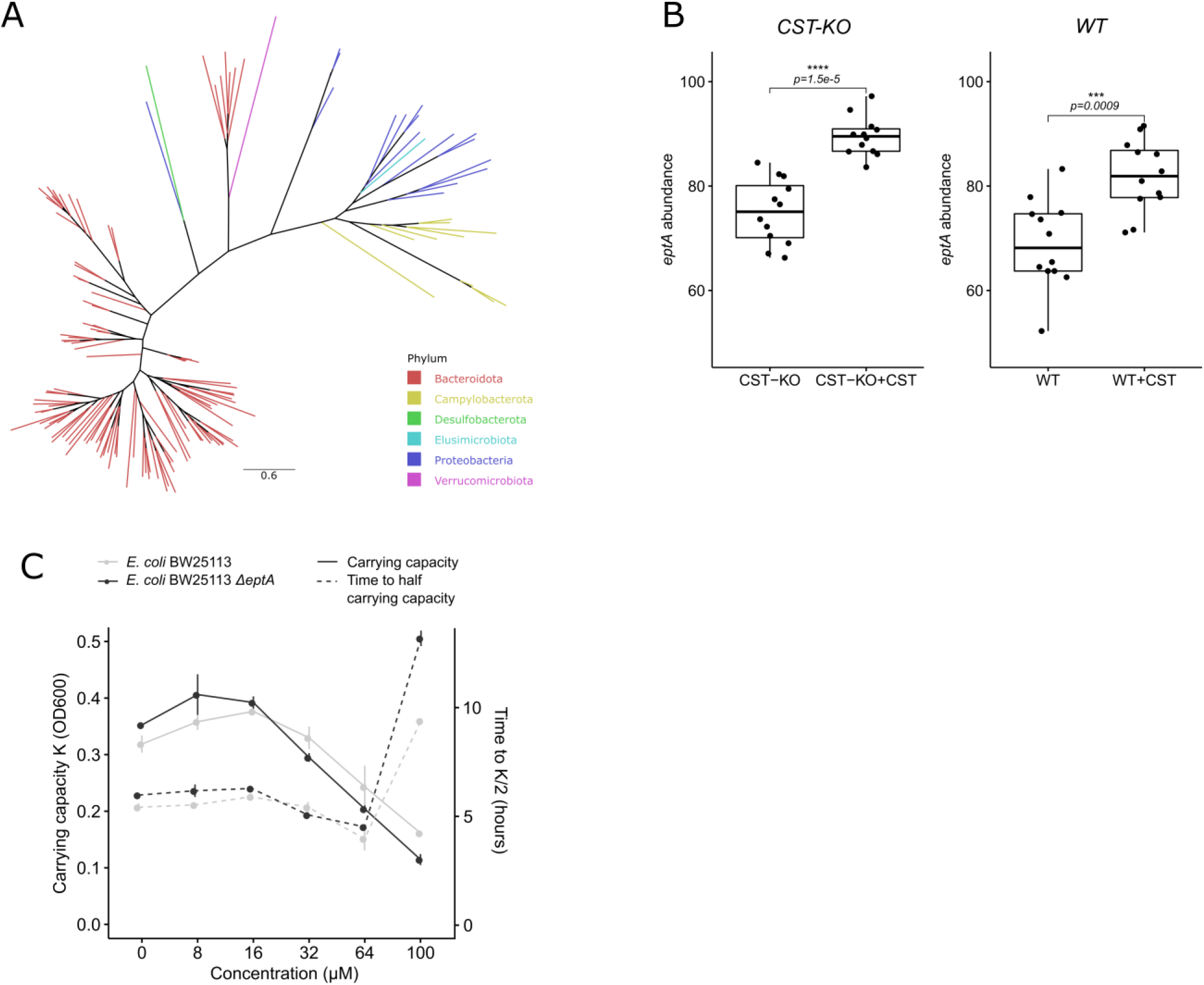
CST treatment promotes the abundance of the antimicrobial resistance gene, *eptA*. (**A**) Unrooted phylogenetic tree depicting the distribution of orthologous *eptA*-like genes throughout bacterial species in the mouse gut. The tree was pruned using Treemmer to aid visualization, while branches were colored by the phylum they belong to. (**B**) Boxplot showing a significant abundance increase of *eptA* harboring bacteria upon CST treatment, both in CST-KO and WT mice. The *eptA* abundance is defined as the sum of abundances from all taxa harboring an *eptA* gene. Boxes represent the median with interquartile range, and whiskers represent the maxima and minima. Significance was assessed by an unpaired *t*-test. (**C**) CST inhibits the growth of wild-type *E. coli* BW25113 and *E. coli* BW25113^*ΔeptA*^ strains, visualized by carrying capacity K (a measure of maximum OD600) and time taken to reach half the carrying capacity (inflection point of the growth curve at mid-log-phase) that depends on CST concentrations ranging from 8-100 μM.

Using the taxonomical distribution of *eptA*, we reanalyzed the abundance of genera harboring *eptA* upon CST treatment. Here, both CST-treated groups showed a significant increase in microbiota-harboring *eptA*, though the effect in the CST-KO group seemed to be more pronounced **(Figure 3B)**. To further support our findings that *eptA*-harboring core taxa increase in abundance in CST treated groups, we looked at individual log-fold changes of single taxa, as described above **(Figure 2)**. Indeed, *eptA*-harboring core taxa were higher in abundance in CST treated groups, regardless of the genotype **(Supplement Figure 3A)**.

To further confirm the bioinformatics analysis outcome for the involvement of the *eptA* gene in CST resistance, wild-type *E. coli* BW25113 and an *eptA* knockout strain (*ΔeptA*) were employed and were cultured in different concentrations (8-100 μM) of CST **(Figure 3C)**. The results indicate that the lack of *eptA* led to a decreased carrying capacity at higher CST concentrations **(Figure 3C)**, which is equivalent to the maximum population size of the culture, as well as a concentration-dependent prolonged lag-phase during the growth of the *E. coli* **(Figure 3C)**.

Similar to the majority of known antimicrobial peptides, CST is recognized by the PhoPQ system, which involves the induction of expression of the gene *pmrD*, followed by activation of *pmrA*, and eventually *eptA* expression ^23^. To this end, we tested the expression of the above-mentioned transcription regulators in *E. coli* BW25113 strain. Most notably, *phoP* showed approximately 3-fold increase at 0.5 h after stimulation of *E. coli* BW25113 strain stimulated with 20 μM CST (sufficient concentration for detection without growth penalty), but only 2-fold and less prolonged in the control, before returning to baseline expression. The *pmrA* showed a later upregulation (after 1h) compared to *phoP* in *E. coli* BW25113 CST-stimulated cultures, while *pmrD* expression spiked in the presence of CST, but was invariant between the different conditions (**Supplementary Figure 3B**).

Taken together, the results show a higher abundance of the bacteria-harbouring *eptA*-like genes upon CST treatment, potentially providing these CST-resistant bacteria an advantage to colonize the gut and, in turn, indirectly affect the colonization of other bacterial taxa.

### Catestatin peptide is degraded by *E. coli* omptin protease

The ability of *E. coli BW25113^ΔeptA^* to still resist comparatively high CST concentrations lead us to test whether the bacterium harbours an additional resistance mechanism, such as degradation or uptake, similar to what we observed in *B. thetaiotaomicron* **(Figure 3D)**. We analyzed the spent culture supernatant by reverse-phase HPLC, which showed absence of CST-corresponding peaks after 24 h of incubation with *E. coli BW25113^ΔeptA^* as well as *E. coli* BW25113 **(Supplementary Figure 4A)**. To test whether the enzyme involved in the CST degradation is excreted in the supernatant, sterile-filtered supernatants of non-stimulated *E. coli* BW25113 cultures were incubated with CST and subjected to tricine SDS-PAGE to visualize the degradation/uptake of CST. Indeed CST was cleaved by *E. coli* BW25113 after incubation for different time intervals from 30 min up till 24 h **(Figure 4A)**. This suggested the involvement of a secreted or membrane-anchored protease, which can also be present in secreted vesicles, in CST degradation. To identify which protease was involved, literature search was performed and revealed that *E. coli* harbours only 2 major outer membrane proteases, predicted to cleave CST ^24^. Among those proteases is the omptin, an outer membrane protease, encoded by *ompT*. To confirm that omptin was responsible for the CST degradation, an *E. coli ompT* knockout strain (*E. coli* BW25113^*ΔompT*^) was employed and was cultured in different concentrations of CST. Compared to the wild-type and *E. coli* BW25113^*ΔeptA*^ strains, *E. coli* BW25113^*ΔompT*^ showed no growth at 100 μM CST and an extensively prolonged lag-phase at 64 μM (**Figure 3D and Figure 4B)**. Collectively, the results imply that wild-type *E. coli* BW25113 possesses the ability to cleave CST through its outer membrane protease omptin, providing the bacterium with another defense mechanism, together with EptA, to resist the antimicrobial action of CST.

**Figure 4.**
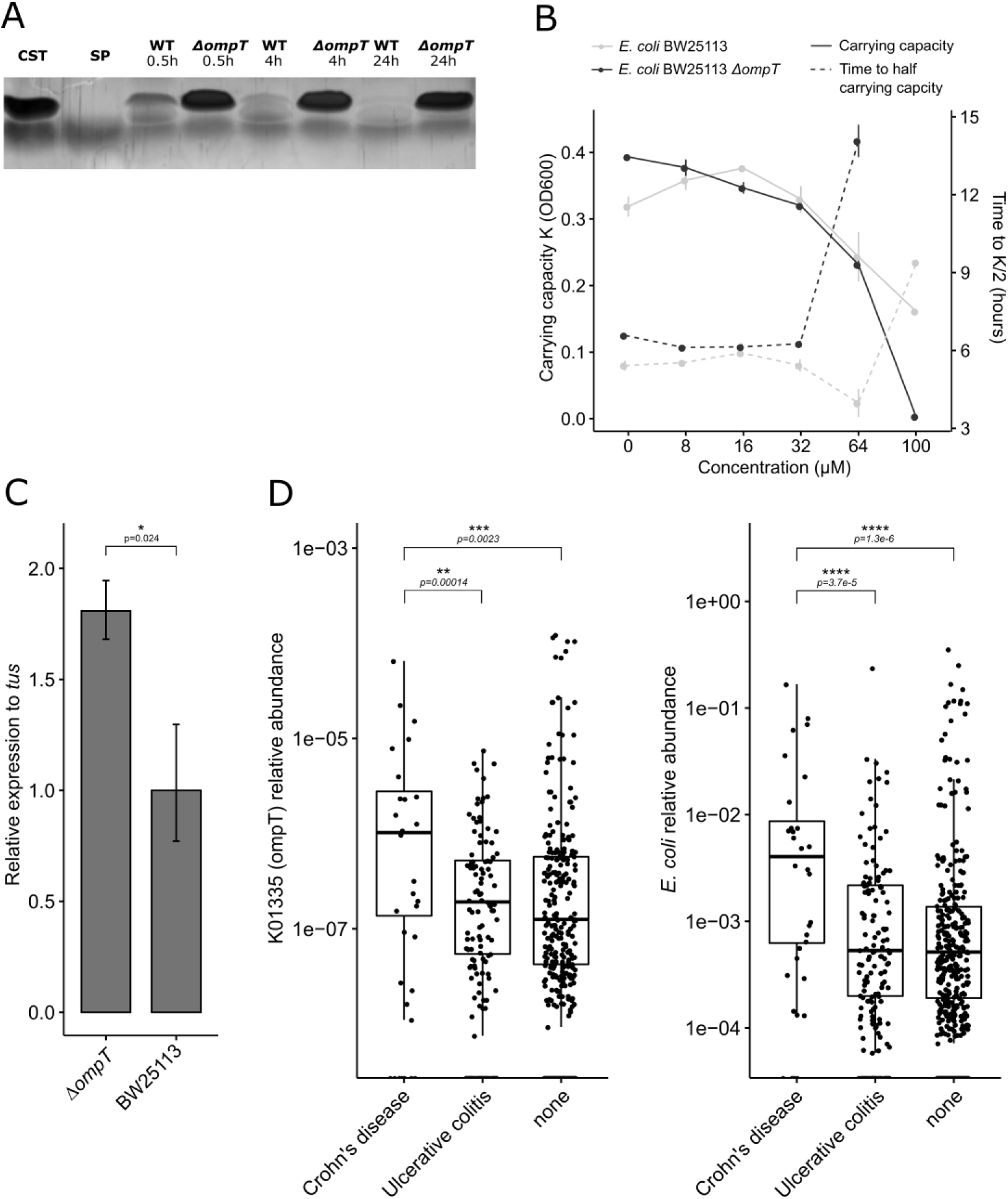
Omptin enzyme is responsible for the degradation of CST in *E coli*. (**A**) Silver-stained tricine SDS-PAGE gels showing the cleavage of CST by wild-type *E. coli* BW25113, but not in *E. coli* BW25113^*ΔompT*^. Sterile non-stimulated culture supernatants were cultured with 100μM CST and samples were collected at different time points; CST: catestatin (2.3kDa), SP: substance P (1.3kDa) (**B**) CST inhibits the growth of *E. coli* BW25113^*ΔompT*^, visualized by carrying capacity K (a measure of maximum OD600) and time taken to reach half the carrying capacity in CST concentrations ranging from 8-100 μM. (**C**) Baseline expression of *eptA* is increased in *E. coli BW25113^ΔompT^* compared and normalized to that of wild-type *E. coli* BW25113. *tus* was used as a housekeeping gene for normalization in both strains and statistical significance was calculated using a Wilcoxon test. (**D**) Relative abundance of omptin like genes or *E. coli* in patients with Crohn’s disease, ulcerative colitis, and healthy subjects. Axes are on log10-scale and significance was assessed using Wilcoxon test without stratification.

To further confirm the complementary function of EptA and omptin, we performed qPCR on control (unstimulated) and CST-stimulated cultures of wild-type *E. coli* BW25113 and *E. coli* BW25113^*ΔompT*^. The expression of *eptA* was upregulated about 1.7-fold in *E. coli BW25113^ΔompT^* without CST stimulation compared to the wild-type strain (**Figure 4C**). When stimulated with 20 μM CST, an increase in the expression of *ompT*, but not *eptA*, was detected *in* wild-type *E. coli* BW25113 without stimulation and at 0.5 h after stimulation, while this effect was reversed at later time points (**Supplementary Figure 3B**).

As *E. coli* BW25113 is considered a lab-strain, which might not be representative of naturally occurring *E. coli* strains, we also tested for the presence of *ompT* in two gut isolate strains, namely *E. coli* Nissle 1917 and *E. coli* DSM 11250. Both were found to be *ompT* carrying (**Supplementary Figure 4B**). Taken together, the results indicate the capacity of *E. coli* strains harbouring omptin to degrade CST, plausibly altering endogenous levels of the peptide in the gut in cases where the abundance of *E. coli* is significantly increased, such as in inflammatory bowel disease ^25^ To test this hypothesis, we investigated the abundance of *ompT*-like genes (KO term K01355) in different subtypes of inflammatory bowel diseases (IBD), which are also associated with altered levels of CST ^15^, using publicly available data from the MetaQuery database ^26^. A significant increase in the abundance of *ompT*-like genes was found between non-IBD and Crohn’s disease patients. Similarly, there was a significant increase in the abundance of *ompT*-like genes between Crohn’s disease patients and patients with ulcerative colitis, but not between ulcerative colitis diseased and non-IBD subjects (**Figure 4D**). This observation was consistent with the abundance of *E. coli* in those patients (**Figure 4D**). Overall, the data indicate that CST degradation by bacteria-harbouring omptin-like proteases may play a role in altered levels of the peptide previously detected in IBD patients, which in turn would result in an overgrowth of these bacteria and worsening of the disease situation.

## Discussion

This study unravels the crucial role of CST on the microbiota composition in the fecal samples of WT and CST-KO mice. Specifically, there was a significant increase in the abundance of taxa that harbor the CST peptide-resistance enzyme, phosphoethanolamine transferase in CST-KO and WT mice treated with CST **(Figure 3B)**. The shifts observed in the microbial phyla in this study after treating WT mice with CST are in agreement with the findings of Rabbi *et al*., who also reported an increase in *Bacteroidota* and a decrease in *Firmicutes*, albeit a different route for the administration of CST and dosage ^19^. Low abundance of Bacteroidota, especially *Bacteroides/Prevotella* have been linked to obesity, and CST-KO mice were described to be insulin resistant and obese besides having elevated blood pressure ^13,27^. In contrast, increased *Bacteroidetes/Firmicutes* ratio has been associated with IBD ^28,29^, which is also characterized by elevated levels of CST ^15^. Nevertheless, as CST is a multifunctional peptide, acting directly on the microbiota as well as on the host, any changes related to CST treatment should be viewed as a complex interplay and not direct causation originating from the microbiota ^12^.

CST treatment could, only to a small extent, restore the microbial composition in a CST-depleted environment, to the composition detected in an environment already primed with CST (WT mice) **(Figure 2)**. In general, the effect of CST treatment observed in the CST-KO group was significantly greater than that in the WT group **(Figure 1)**. This difference could be attributed to the presence of normal levels of CST in the WT mice; thus, the microbiota is already primed with the peptide and has evolved mechanisms to resist its antimicrobial effect. In contrast, the CST-KO group, in which CST is absent, may harbor bacteria that are susceptible to the CST antimicrobial activity but were eradicated upon CST treatment **(Figure 2)**. Especially the core taxa resisting CST treatment seem to benefit from their presence in the gut lumen environment, by exhibiting an increase in abundance. Introduction of CST to a CST-depleted environment did not necessarily promote exclusively health beneficial microbiota, but rather led to a more equilibrated composition, including also pathobiont taxa. For example, in CST-treated CST-KO, *Oscillibacter* and *Mucispirillum* **(Figure 2)**, both of which increased in relative abundance, have been found to increase in diet-induced obesity and might be considered as pathobionts ^30,31^. Similarly, *Alistipes* was shown to have a detrimental influence on colorectal cancer progression, despite contrasting evidence that it also has protective effects against colitis or cardiovascular diseases ^32^. On the other side *Odoribacter*, a SCFA-producer, which increased in relative abundance in CST-treated WT mice **(Figure 2)** has been linked to immunomodulatory effects and general health benefits presumably also in cooperation with *Akkermansia* ^33^.

So far, antimicrobial resistance mechanisms have been mostly investigated in pathogens due to their implication on antibiotic resistance. In *in-vitro* studies, CST has been reported to have an inhibitory effect against the pathogenic gram-positive (*Micrococcus luteus*, *Bacillus megaterium*), gram-negative bacteria (*Escherichia coli* D22), and filamentous fungi (*Neurospora crassa and Aspergillus fumigatus*) ^5^. Moreover, human CST showed antimicrobial effects against several skin microbes including the gram-positive (*Staphylococcus aureus* and Group A *Streptococcus*), gram-negative bacteria (*E. coli* O29, and *Pseudomonas aeruginosa*), also (*Candida albicans*), and filamentous fungi (*Aspergillus niger, Aspergillus fumigatus, Trichophyton rubrum*) ^16^.

As a defense mechanism, bacteria have evolved several systems to survive the effect of the antimicrobial peptides ^21^. The ability to withstand these peptides provides a potential benefit in colonization over other members of the gut microbiota especially under pathological conditions ^34^. Analogously, antimicrobial peptides are regularly used by the host to select for specific gut microbiota in homeostatic situations. For CST, the antimicrobial activity resides in the N-terminal of the peptide, which is highly cationic ^5^. These positive charges in the peptide interact with the anionic components of the bacterial cell membrane, resulting in the permeabilization of microbial membranes, leading to cell lysis ^7^. The present study suggests that CST may as well play a role in aiding the host to select for which microbes to colonize the gut. The abundance of bacteria encoding the phosphoethanolamine transferase genes increased upon CST treatment **(Figure 3A-E)**. This is not surprising since the *E.coli* phosphoethanolamine transferase e*ptA*, for example, has been linked to colistin resistance, being a cyclic proteinogenic antibiotic ^35^. EptA acts by modifying core lipopolysaccharide components with ethanolamine, increasing membrane charge and thus, repelling the antimicrobial agent. The fact that EptA-like proteins are abundant across a variety of different taxa present in the human gastrointestinal tract **(Figure 3A-C)** implies that antimicrobial resistance is a mutualistic and important component in the selection of a specific microbiota composition by the host, through active modulation of antimicrobial peptide levels, such as CST. Indeed, this paradigm is supported by our findings that CST treatment induced a large-scale alteration in the microbiota composition not only in the CST-KO group but also in the WT group, both of which clustered closer to each other, and further away from the untreated groups **(Figure 1)**.

The induced expression of *eptA* gene in *E. coli BW25113^ΔompT^* **(Figure 4C)** implies a complementary mechanism when other defense mechanisms are missing. Indeed, we uncovered the degradation of CST by wild-type *E. coli*, a member of the gut microbiota via omptin, an outer membrane protease, known to cleave other cationic antimicrobial peptides, such as protamine or LL-37 ^36,37^. The presence of these fragments with high proteolytic activity in the gut has its implications on the microbiota composition, as it may facilitate colonization of other CST-susceptible bacteria ^38^. It remains elusive though, whether CST is taken up and accumulating inside the bacteria or is further degraded, and what the fate of the resulting cleavage fragments is, e.g., through uptake by other microbiota or interaction with the host. Analogously it remains unclear how other bacteria, such as *B.thetathiomicron* can degrade CST **(Supplementary figure 2)** and how widespread such a phenomenon is among the gut bacteria.

Levels of CST are increased in IBD patients, while *E. coli* was associated with the development and progression of different IBD types ^25,39^. The higher abundance of *ompT*-like genes in samples from Crohn’s disease patients, which coincided with a higher abundance of *E.coli* **(Figure 4D)**, suggests that bacteria harboring *ompT*-like genes, such as *E. coli*, may have a benefit to colonize the gut due to its ability to resist CST. In cases where the overall microbial composition is compromised, and the endogenous levels of CST are altered, like in IBD, such bacterial resistance may cause particular bacterial strains to be involved in worsening the gut inflammation. Overgrowth of gut bacteria with the capacity to resist CST may lead the host to secrete higher levels of CST as a defense mechanism. The levels of CST were reported to be higher in IBD patients compared to healthy subjects ^15^. It remains unclear why only samples from Crohn’s disease patients but not ulcerative colitis showed a higher abundance of *ompT* as well as *E. coli*. One plausible explanation could be due to the different *E. coli* strains predominantly present in either Crohn’s disease or ulcerative colitis ^25^ **(Figure 4D)**.

Overall, the present study highlights the significant role of the CST peptide in determining the microbiota composition. The fact that gut bacteria with a capacity to resist the antimicrobial effect of CST become more abundant upon CST treatment, suggests that these bacteria may have gained an advantage over other microbiota members to survive the gut environment, which, indirectly shapes the colonization of the total community. Further studies should also explore the changes elicited by CST administration on host physiology, which might in turn affect the gut microbiota composition. One limitation of the present study is the relatively uncontrolled actual concentrations of CST that eventually reach the gut after intraperitoneal injection. This concentration variability could increase interindividual variability in the microbial composition.

## Materials and Methods

### Animals

All studies with mice were approved by the University of California San Diego (UCSD) and Veteran Affairs San Diego Healthcare System (VASDHS) Institutional Animal Care and Use Committees for the Mahata laboratory (UCSD: #S00048M; VA: #A13-002) and were performed in San Diego in adherence to the NIH Guide for the Care and Use of Laboratory Animals. Our experiment was based on a single cohort of mice which constisted of twelve mice per group each (control and CST-treatment, per genotype); out of those, twenty four male adult C57BL/6 J mice (age 20 weeks) were purchased from Jackson Laboratory (Bar Harbor, ME) and twenty four CST-KO mice (age 20-28 weeks) generated in the Mahata laboratory, were used. CST-KO mice have a deletion in the 63 bp CST domain from Exon VII of the *Chga* gene ^27^ in C57BL/6 background. Mice from WT and CST-KO groups were injected intraperitoneally once daily with CST (2 μg/g body weight; Genscript Biotech Corporation, Piscataway, NJ) for 15 days based on previous experimentation ^15^. Mice were housed in 4 to 5 animals per cage and had free access to water and food (Normal Chow Diet, LabDiet 5001) in temperature and humidity-controlled rooms with a 12-hr light/dark cycle.

### Determination of short-chain fatty acids (SCFAs) in cecal samples

Cecal samples (100 mg) were suspended in 1 mL of saturated NaCl (36%) solution. An internal standard (50 μL of 10.7 μM 2-methyl butyric acid in MQ water) was added and the samples were homogenized using glass beads. After the addition of 150 μL H_2_SO_4_ 96%, SCFAs were extracted with 3 ml of ether. The ether layer was collected and dried with Na_2_SO_4_ (150 mg). The supernatant (0.5 μL) was analyzed using gas chromatography with flame ionization detection (Agilent, Santa Clara, California, USA). The system was equipped with a DB FFAP analytical column (30m x 0.53 mm ID, 1.0 μm; Agilent) and helium GC grade (5.6) was used as carrier gas with a constant flow of 4.2 ml/min. The initial oven temperature was held at 100 °C for 3 min, ramped with 4 °C/min to 140 °C (isothermal for 5 min) and further with 40 °C/min to 235 °C (isothermal for 15 min). The resulting chromatograms were processed using ChemStation (Agilent Technologies).

### DNA isolation and 16S rRNA gene sequencing

DNA isolation was performed on fecal samples that were collected from untreated and CST-treated WT and CST-KO mice. A phenol-chloroform-isoamyl alcohol procedure was used for DNA extraction ^40^. Briefly, the pellet was resuspended in 1 ml of lysis buffer (940 μl TE buffer, 50 μl SDS 10% and 10 μl Proteinase K 20 mg/ml) in a 2 ml screw cap micro-tube containing a mix of zirconium and glass beads. Then, samples were incubated at 58 °C for 1 hour and 150 μl of buffered phenol (Invitrogen, 15513-047) was added. To support lysis, samples were homogenized 3 × 30 sec with 1-min intervals on ice in a mini bead-beater (Biospec, Bartlesville, USA). This was followed by the addition of 150 μl chloroform/isoamyl alcohol (24:1) and centrifugation at 16,000x *g* for 10 min and 4 °C. The upper layer was carefully transferred to a clean tube, 300 μl of phenol-chloroform-isoamyl alcohol [25:24:1] was added, centrifuged and the process was repeated by adding 300 μl of chloroform-isoamyl alcohol [24:1]. To precipitate the DNA, the upper layer was transferred to a new tube and 1 volume of absolute isopropanol and 1/10 volume of 3 M sodium acetate was added. Samples were incubated overnight at −20 °C. Subsequently, samples were centrifuged at 16,000x *g* for 20 min at 4 °C, the supernatant was discarded and the pellet was washed with 700 μl of 70% ethanol. After ethanol aspiration the DNA pellet was air-dried for 30 min and resuspended in 100 μl TE buffer. In each step, samples were vigorously mixed using a vortex.

Sequencing of the V3-V4 region of the bacterial 16S rRNA gene was carried out by Novogene Co. Ltd. Briefly, for sequencing library preparation, raw DNA extracts were diluted to 1 ng/μl in sterile water, and amplicons were generated by PCR (primers 341F and 806R) using a Phusion High-Fidelity PCR Master Mix (New England Biolabs). Amplification product quality was assessed by gel electrophoreses and samples were pooled in equimolar ratios. Libraries were generated with a NEBNext Ultra DNA Library Prep Kit for Illumina and sequencing was carried out on an Illumina MiSeq 250 bp paired-end platform. Initial processing of reads involved trimming of adapters and primers using a Novogene in-house pipeline (Novogene Co. Ltd, Cambridge, UK).

### Microbiota analysis

Paired-end sequencing reads were filtered, denoised, merged, and classified with the *dada2* package in the statistical programming language R while processing forward and reverse reads separately until merge ^41,42^. Briefly, reads were truncated to 220 bases and low-quality reads were filtered followed by dereplication. Error models were learned, while manually enforcing the monotonicity of the error function. Reads were denoised and merged with minimal overlap of 12 bp, while non-merging reads were concatenated. Singletons have been removed before performing bimera removal, followed by read classification using SILVA (V138) as a taxonomical reference database.

For downstream analysis, the *phyloseq* and *microbiome* packages were used, samples were rarefied to even depth and richness was determined by the number of observed amplicon sequence variants (ASVs), as well as ACE index and alpha diversity was assessed via Shannon’s *H* and inverted Simpson indexes ^43,44^. All ASVs were collapsed on genus level and cumulative sum scaling was applied using *metagenomeSeq* ^45^. The resulting genus abundance table served as input for different types of ordinations such as principal component analysis (PCA) or redundancy analysis (RDA). The significance of constraints in constrained RDA was determined by permutation-like ANOVA. Differential abundance was assessed by unpaired Wilcoxon Rank Sum test using *p* < 0.05 as a significance threshold followed by FDR correction.

To generate the double log-fold change plot, first, the core microbiota were estimated for each group by filtering raw ASV counts to contain only more than 10 reads and be present in more than 10% of the samples. Next, only taxa intersecting between CST treated groups were selected and analyzed further. Differential abundance was assessed by a Wilcoxon Rank Sum test between treated and untreated groups per genotype and FDR corrected. Only taxa with an adjusted *p* < 0.05 were considered further. Their mean log-fold changes between groups were calculated and plotted against each other.

Further LefSe analysis was carried out, adjusting the alpha value for factorial Kruskal-Wallis test and the alpha value for pairwise Wilcoxon tests to 0.01. Additionally, the LDA score threshold was set to 3.0 ^46^

For the functional analysis, the metagenomic prediction from 16S rRNA gene sequencing data was done using PiCRUSt2, using raw ASVs and unnormalized ASV abundances ^47^. For this, ASVs were first filtered to have a minimal abundance of 10 reads and to be present in at least 1/10 of the samples. PiCRUSt2 was run with standard parameters and stratification enabled. The metagenomic data was further analyzed by using STAMP v2.1.3 ^48^. For statistical analysis, White’s non-parametric t-test, two-sided, and Benjamini-Hochberg multiple test correction were used to reveal differences between groups.

### Phylogenetic analysis

The amino acid sequence of the *E. coli eptA* gene (UniProt P30845) was searched with *phmmer* from the hmmer suite (v3.3.1; hmmer.org) against a non-redundant dataset of the mouse gastrointestinal bacterial catalog (MGBC) ^49^. Hits were filtered on full sequence E-value, excluding all sequences with an E-value greater than 0.5x the median absolute deviation from the median. Additionally, only sequences originating from genomes with >90% completeness and <1 contamination were taken into account. Sequences were dereplicated, aligned with MAFFT (v7.453) and a phylogenetic tree was constructed using FastTree (v2.1.11) ^50,51^. The produced gene tree was visualized with FigTree (v1.4.4), to aid visibility the tree was pruned using Treemmer (v0.3; with parameter -RTL 0.95) ^52^

### Bacteria and growth inhibition assay

*E. coli* BW25113, BW25113^*ΔompT*^ (JW0554) and BW25113^*ΔeptA*^ (JW5730) (**Supplementary Table 1**) were routinely grown aerobically in Difco Antibiotic Medium 3 (BD, Franklin Lakes, USA) at 37 °C degrees without agitation. Bacteria were inoculated from −80 °C stocks and grown overnight. Before the experiment, cultures were diluted at 1:100 in fresh medium from overnight cultures. Five different CST concentrations were tested in triplicates in a 96-well plate. For this, 10 μl of a 10x concentrated stock of CST were added to 90μl of diluted culture to obtain final concentrations of 8, 16, 32, 64, and 100 μM. Bacterial growth was monitored for 24 h at 600 nm (OD600) with shaking for 5s every 15 min using a BioTek ELx808 microplate reader (BioTek, Winooski, USA). Growth curves were fit using the *growthcurver* R package (v0.3.0).

### Tricine SDS-PAGE

Overnight cultures of *E. coli* strains BW25113 and BW25113^*ΔompT*^ **(Supplementary Table x)** were centrifuged at 10,000 g for 10 min at 4°C and the supernatant was collected. The supernatant was further sterile filtered (0.2 μm filters) and to 90 μl of supernatant 10 μl of a 10x CST stock was added to reach a final concentration of 100 μM. Aliquots of 20 μl were taken at time points 0.5, 4, and 24 h.

To visualize CST degradation, culture supernatants were run on Tricine SDS-PAGE following a modified protocol ^53^. Briefly, gels consisted out of a 4% stacking gel and a 16% resolving gel both with a cross-linker ratio of 19:1, and were prepared to standard procedure. For sample prep, 10 μl of each sample were mixed with 10 ml of reducing sample buffer (100 mM Tris-HCl, 1% SDS, 4% 2-mercaptoethanol, 0.02% Coomassie Brilliant Blue, 24% glycerol) and incubated at 70°C for 5 min. Subsequently, 10 μl of the sample was loaded on the gel, first, run for ~30 min at 30 V, and then at 170 V for 90 min. Silver staining was applied to visualize proteins using various modifications in fixation to avoid loss of small proteins ^54^. For this, gels were fixed (30% ethanol, 15% formalin, 5% acetic acid) for 60 min and quickly washed once with 50% ethanol and twice with MilliQ-filtered water. Gels were incubated for 40 min in 0.005% sodium thiosulfate, followed by incubation for 40 min in 0.1% silver nitrate solution containing 0.07% formalin. Washing with MilliQ-filtered water was performed and a 2% sodium carbonate solution containing 0.1% formalin was added. Gels were incubated for 1-2 min until bands became visible and development was stopped by removing the developer and adding 50 mM EDTA solution. Gels were then imaged in a ChemiDoc MP Imaging System (BioRad, Hercules, USA).

### Expression analysis

Overnight cultures of *E. coli* BW25113, *E. coli* BW25113^*ΔeptA*^, and *E. coli* BW25113^*ΔompT*^ were resuspended 1:1000 in 6 ml of fresh Difco Antibiotic Medium 3 and grown until OD600 0.5 was reached. Aliquots of 1 ml were harvested, centrifuged at 10,000 g for 10 min, and immediately stored at −80 °C. This and all subsequent centrifugation steps were done at 4 °C.

For culture stimulation with CST *E. coli* BW25113 or *E. coli* BW25113^*ΔompT*^ was resuspended 1:1000 in 6 ml fresh Difco Antibiotic Medium 3 and grown until OD600 0.5. An aliquot of 5 ml was harvested and centrifuged at 3,000 g for 10 min. The pellet was resuspended in fresh medium containing 20 μM CST and gently mixed. Samples of 1 ml were taken at time points 0h (before resuspension), 0.5h, 1h, 2h, and 4h, subsequently centrifuged at 10,000 g for 10 min and immediately stored at −80 °C.

RNA was isolated using a phenol:chloroform:isoamyl alcohol procedure as previously described ^55^. The harvested bacterial pellet was resuspended in 200 μl TE buffer and transferred to a tube containing 0.5 g of 0.1 mm zirconia/silica beads, 25 μl 10% SDS, and 200 μl phenol:chloroform:isoamyl alcohol solution. The subsequent procedure was performed as described and isolates were stored at −80°C until further use.

qPCR was carried out according to standard procedures ^56^. Briefly, 2 μg of RNA were reverse transcribed using a High-Capacity cDNA Reverse Transcription Kit with RNAse Inhibitor (Applied Biosystems, Vilnius, Lithuania). Subsequently, 10 ng of cDNA were used for qPCR employing a PowerUp SYBR Green Master Mix (Applied Biosystems, Vilnius, Lithuania) and primers for genes *eptA, ompT, phoP, pmrD, pmrA* and *tus*. Reactions were performed in a clear 96-well plate using a CFX96 Real-Time PCR System (BioRad, Hercules, USA). Gene expression was assessed by normalizing to housekeeping gene *tus* using the R package *qpcR* (v1.4) for curve fitting and the *pcr* (v1.2.2) package for differential expression calculation employing the 2^−ΔΔCt^ method as described in ^57^.

### Statistical analysis

All statistical tests were performed using GraphPad Prism 7 or the statistical programming language R. Specific statistical tests are indicated in the text or figure legends. Normality was assessed by either D’Augustino-Pearson omnibus normality test or Shapiro-Wilk normality test. If normality was met a t-test was chosen, otherwise non-parametric Mann-Whitney was used. Outliers were assessed with GraphPads ROUT method (Q=1).

### Data availability statement

All data generated or analyzed during this study are included in this manuscript and its supplementary information files. The 16S rRNA gene amplicon sequence data were deposited under BioProject number PRJNA741992.

## Authors’ Contributions

P.G-D., M.S, S.E.A and S.K.M conceived and designed the study. P.G-D., M.S., B.D, and S.K.M. performed the experiments, and P.G-D., M.S, B.D., and S.E.A analyzed the data. P.G-D, M.S, S.E.A. wrote the original manuscript that was reviewed by A.D, B.D., K.V., and S.K.M. Funding for these studies were acquired by S.E.A. and S.K.M. All authors read and approved the final manuscript.

## Acknowledgment

P.G-D thanks the National Council of Science and Technology in Mexico (CONACyt) for the Ph.D. grant assigned to CVU 690069. We thank Dr. Matthias Heinemann of Department of Molecular Systems Biology, University of Groningen, the Netherlands, for providing us *ΔeptA* and *ΔompT E. coli* BW25113 mutant strain; Dr. Greet Vandermeulen of Department of chronic diseases and metabolism, Faculty of Medicine, KU Leuven, Belgium for the help with SCFAs analysis.

