## Supplementary information for "Catestatin selects for the colonization of antimicrobial-resistant gut bacterial communities" (from another version of this preprint: version 1, 2021-10-11)

**Title**

[#] Shared first author

Sahar El Aidy,

### Supplementary Methods

#### Culturing of fecal content

A pre-culture was prepared by pooling the fecal content of 20 healthy human subjects, which was previously collected and stored in (1:1) liquid amies at -80 °C. For this, approximately 1.1 g of fecal content from each subject were used and suspended with a total of 71 ml of Gifu Anaerobic Broth (HiMedia Laboratories GmbH, Einhausen, Germany). The culture was gently homogenized by vortexing and transferred into a 100 ml Erlenmeyer flask with a cotton stopper and incubated anaerobically (2% $H_2$, 5% $CO_2$, balanced with $N_2$) in a Coy Laboratory Anaerobic Chamber (neo-Lab Migge GmbH, Heidelberg, Germany) at 37 °C overnight. A 10x stock solution of CST was prepared by dilution with MilliQ-filtered water. For each culture 2.7 ml of pre-culture were added to a tube (5ml tubes; Geiner Bio-One, Frickenhausen, Germany), incubated for 30 min before the addition of 300µl of 10x peptide stock solution, to reach a final volume of 3ml and final concentration of 10 µM CST. Samples of 200 µl were taken after 24 and 48 hours centrifuged at 16000 x g for 15 min, and supernatant and pellet were stored separately at -80 °C until further use.

DNA isolation followed a previously proposed protocol by Yu and Morrison (2004). Briefly, the pellet was resuspended in 750 µl of lysis buffer (500 mM NaCl, 50 mM Tris-HCl, 50 mM EDTA, 4 % SDS) and transferred to a screw cap tube containing ~ 500mg of 0.1mm zirconium bead and four 3mm glass beads. Samples were homogenized 3 × 1 min with 1-minute intervals on ice in a mini bead-beater (Biospec, Bartlesville, USA). Subsequently, samples were further incubated at 95°C for 15min and centrifuged at 16000 x g for 15 min at 4 °C. The supernatant was transferred to a clean tube and 20µl of 10M ammonium acetate was added and incubated on ice for 10 min before centrifuging at 16000 x g for 15 min at 4 °C. The supernatant was again transferred to a clean tube and one volume of 100% ice-cold isopropanol was added. Samples were incubated on ice for 1.5-2 h and centrifuged at 16000 x g for 15 min at 4 °C. The supernatant was aspirated and the pellet was washed with 200 µl of 70% ethanol with the following centrifugation to collect the pellet. Ethanol was removed and tubes were left to air-dry for ~1 h before resuspension of the pellet in 200 µl of TE buffer. Sequencing and microbiome analysis was carried out as described in the main text.

**Inhibition assay**

We determined minimal inhibiting concentrations for several gut-related bacteria, which showed up as differentially abundant taxa in previous unpublished analyses and were thought to be related to a CST environment. Bacteria were grown overnight in 1ml of modified DSMZ medium 110 to stationary phase and subsequently diluted 1/100 in medium. Five different CST concentrations were tested in triplicates in a 96-well plate. For this 10 µl of a 10x concentrated stock of CST were added to 90µl of diluted 6 culture to obtain final concentrations of 4, 16, 32, and 64 µM. Growth was monitored for 24h at 600nm (OD600) with shaking for 5s every 15min using an Infinity 200 Pro microplate reader (Tecan, Grödig, Austria). Growth curves were fit using the growthcurver R package and minimum inhibiting concentrations (MIC) were determined upon visual inspection of the growth curves.

**HPLC**

Bacterial cultures were centrifuged at 16000 x g for 10 min at 4 °C and 100 µl of supernatant were transferred to a clean tube. 200 µl of denaturation solution (7 M urea, 2 M thiourea, 20 mM dithiothreitol) were added, briefly vortexed and 1ml of ice-cold 100% acetone was added. Samples were incubated on ice for 1.5 h before centrifugation at 16000 x g for 15 min at 4°C. The supernatant was discarded and 200 µl of resolubilization solution (70% acetonitrile, 12 mM HCl) was added followed by incubation on ice for 1.5 h. Samples were centrifuged again at 16000 x g for 15 min at 4 °C, the supernatant was transferred to a clean tube and dried in a Savant speed-vacuum dryer (SPD131, Fisher Scientific, Landsmeer, The Netherlands) at 45 °C for ~1 h. The dry pellet was reconstituted in 120 µl of 0.1% trifluoroacetic acid (TFA) and filtered through a 0.2 µm filter (adapted from[2]). HPLC analysis was carried out on a Dionex UltiMate3000 system (Thermo Fisher Scientific, The Netherlands) with a Kinetex C18 reverse-phase column (5 µm, 100 Å, 250x4.6 mm, Phenomenex, Utrecht, The Netherlands) and gradient elution. The mobile phase system consisted of 0.1% TFA/water (solvent A) and 70% ACN/water + 0.1% TFA (solvent B), while the gradient was as follows: 0-10 min 0%B, 10-50 min to 70%B, 50-55 min 70%B. Detection was performed via a Waters 474 fluorescent detector (Waters Chromatography B.V, Etten-Leur, The Netherlands) with excitation wavelength set to 274 nm and emission set to 303nm (slit width 18 nm).

### Supplementary Results

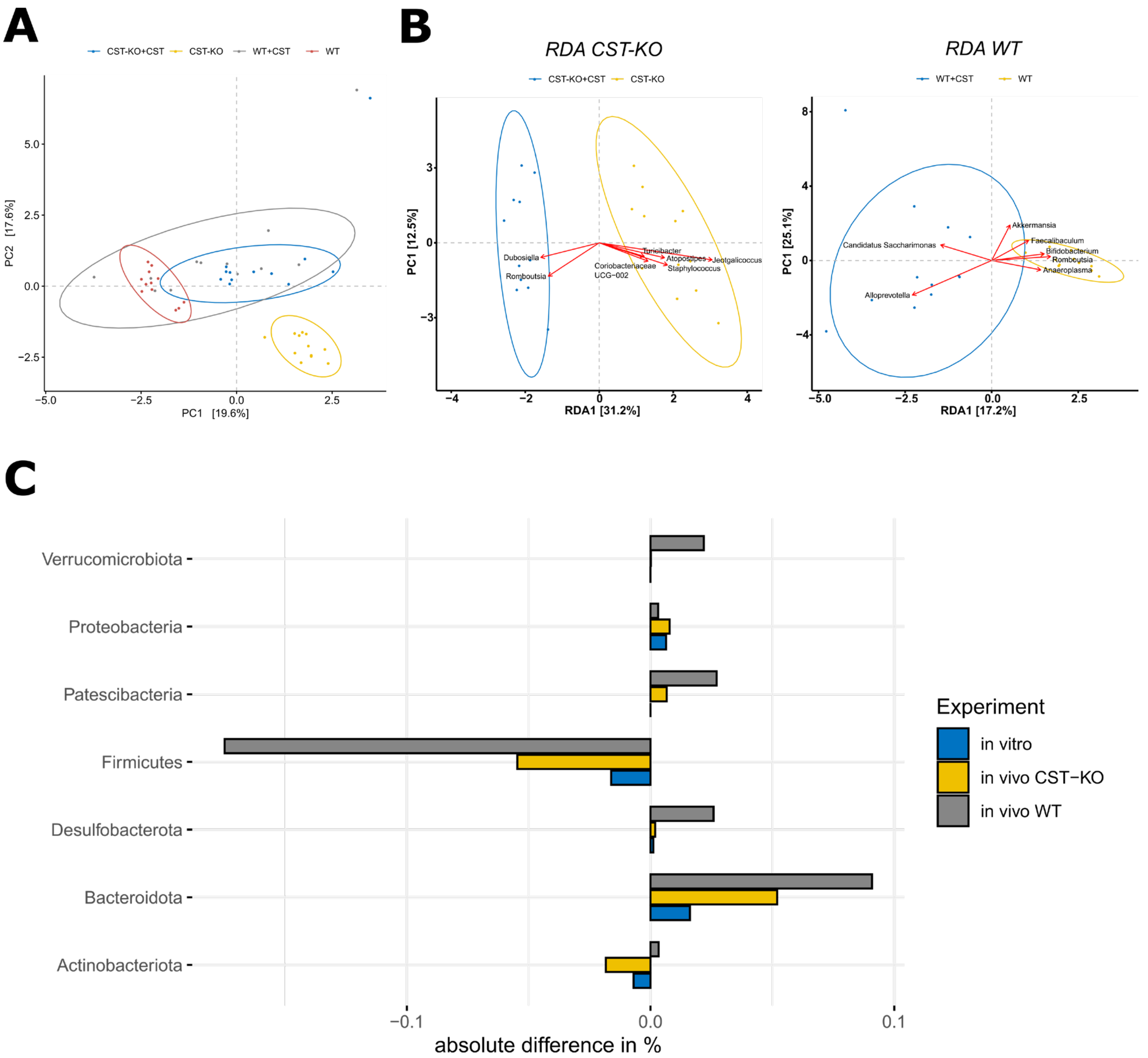


**Supplementary Figure 1. (A)**Principal component analysis (PCA) at the genus level using CSS scaled abundances determined by 16S genotyping. The plot reveals a distinct clustering of treatment groups, with the treated wild-type cluster being greatly spread. Ellipses represent normal data ellipses produced by methods from the R package ggplot2. (**B**) Treatment-constrained RDA of either CST-KO or WT. Arrows indicate taxa associated with either CST-treated (blue) or non-treated (yellow) groups, while the length of the arrow indicates the strength of the association. Ellipses represent normal data ellipses produced by methods from the R package ggplot2. (**C**) Differences in relative microbial compositions on phylum level between CST treated and untreated for the *in vivo* experiment (WT and CST-KO) and non-peptide control and CST for the *in vitro* experiment. Values are expressed as absolute difference, such that 0.1 translates to a relative compositional change of 10%, while only phyla with a mean abundance >1% are displayed for convenience.

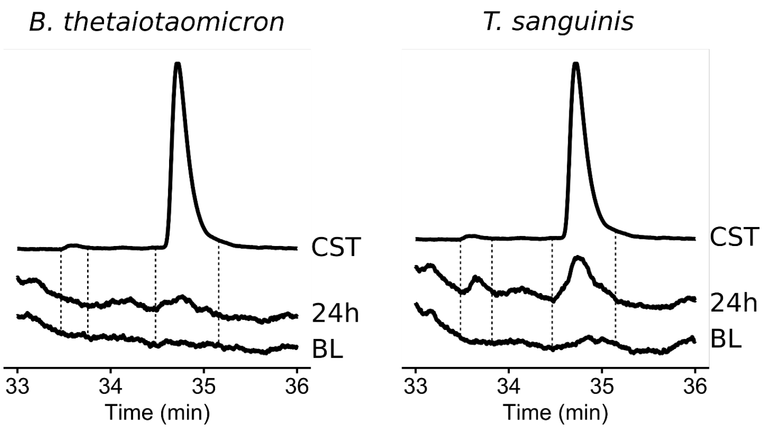


**Supplementary Figure 2**. Reverse-phase HPLC chromatograms of culture for *B. thetaiotaomicron* and *T. sanguinis* stimulated with 32μM CST. CST –catestatin standard, 24h – culture supernatant after 24h of incubation, BL – baseline of culture supernatant without CST.

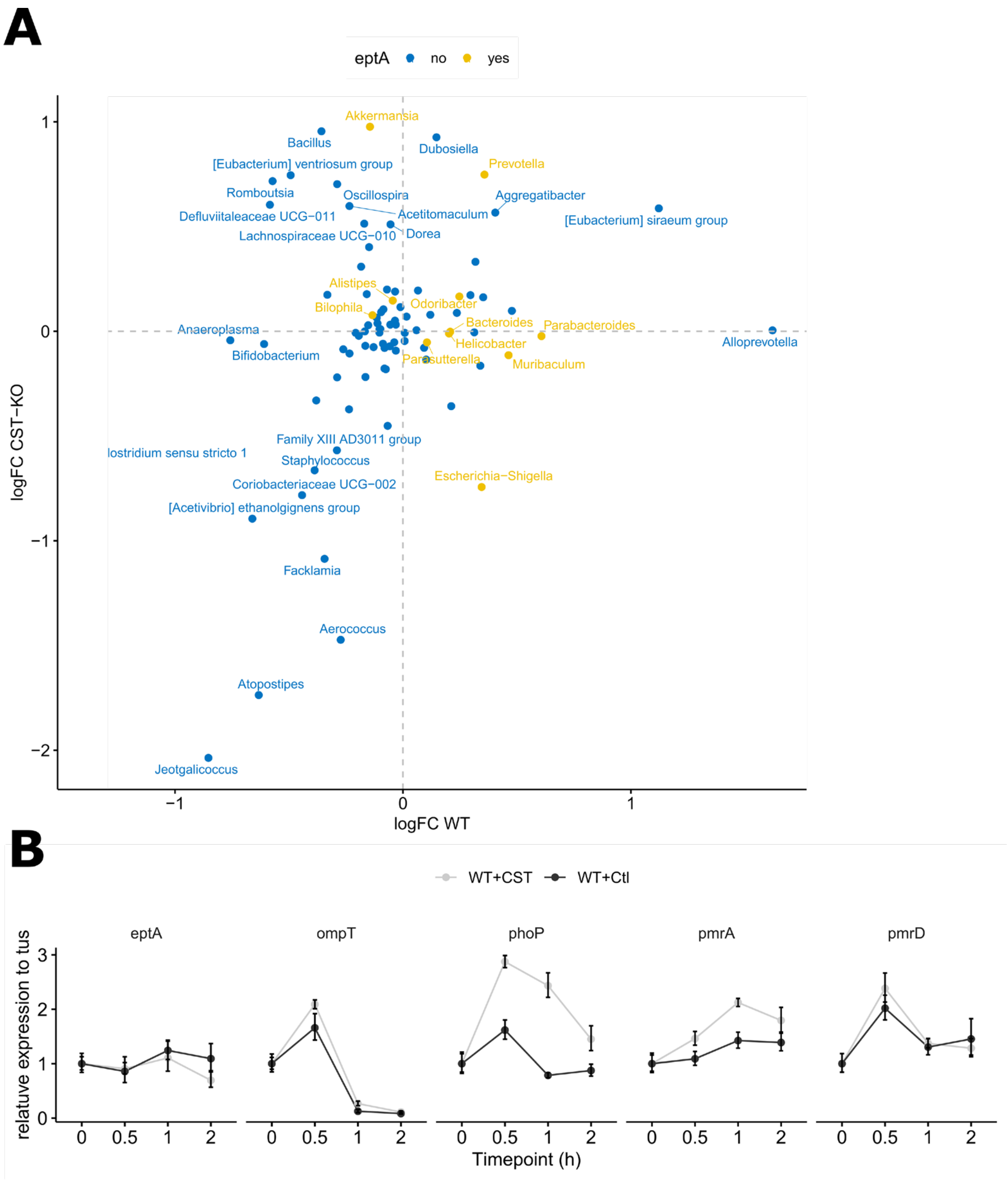


**Supplementary Figure 3. (A)** Scatter plot of log-fold changes, comparing abundance changes for significantly abundant genera between treated and untreated groups per genotype. The coloring indicates the presence of an *eptA*-like gene. (**B**) Expression analysis of wildtype *E. coli* BW25113 stimulated with CST or medium as control (Ctl). *phoP* and *pmrA* are substantially upregulated compared to WT control and knockout, while the increase in *ompT* expression is only minor compared to WT control. *eptA* expression remains invariant upon stimulation. Expression is assessed by the -ΔΔCt method with normalization to housekeeping gene *tus* and timepoint 0 preceding stimulation. Bars indicate point ranges.

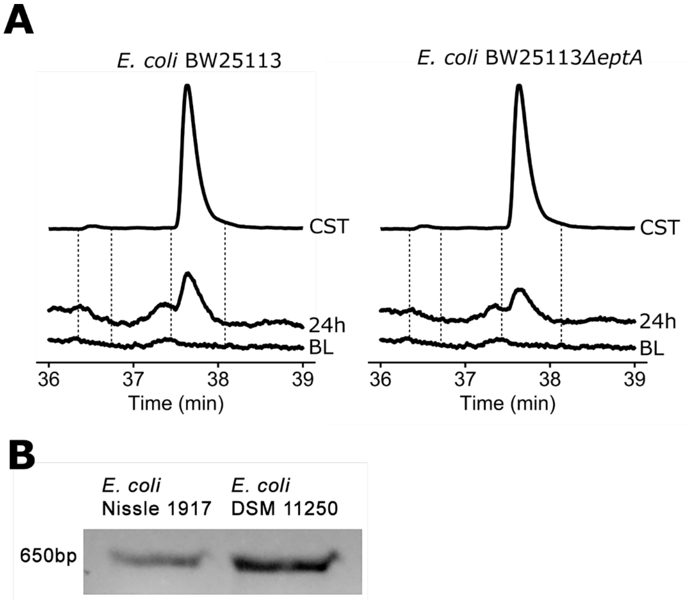


**Supplementary Figure 4**. (**A**) Reverse-phase HPLC chromatograms of culture supernatant from *E. coli* BW25113 and *E. coli* BW25113$^{\Delta eptA}$ stimulated with 32μM CST. CST – catestatin standard, 24h – culture supernatant after 24h of incubation, BL – baseline of culture supernatant without CST. (**B**) Polymerase chain reaction (PCR) of a 650bp fragment of the *E. coli ompT* gene, resolved on 1% agarose for two gut isolates.

**Supplementary Tables**

**Supplementary Table 1.** Resources

| Strain or Primer | | Relevant feature | Reference |
|---|---|---|---|
| Strains | | | |
| | *E. coli* BW25113 | Δ(araD-araB)567, ΔlacZ4787(::rrnB-3), λ-, rph-1, Δ(rhaD-rhaB)568, hsdR514 | [3] |
| | *E. coli* JW5730 | *E. coli* BW25113 with ΔeptA740::kan | [3] |
| | *E. coli* JW0554 | *E. coli* BW25113 with ΔompT774::kan | [3] |
| | *Bacteroides thetaiotaomicron* ATCC29148 | | ATCC |
| | *Turicibacter sanguinis* DSM14220 | | DSMZ |
| Primers | | | |
| | phoP_fwd | GCAGACGTCCCATCAGTACAT | This study |
| | phoP_rev | GTCGATGACGCAGAAGATGC | This study |
| | eptA_fwd | CTCTTCAAGCCCGTGGAACA | This study |
| | eptA_rev | CGCTGATGACACCGCAAATG | This study |
| | ompT_fwd | CGATTCCATGCGCCTTCAAC | This study |
| | ompT_rev | GGAGGCCGAAAAGTCAGTCA | This study |
| | pmrD_fwd | TAACTGCTTGCCGAGAGGAC | This study |
| | pmrD_rev | CTTATGCTCTGCGATGCTGG | This study |
| | pmrA_fwd | ATGCACTTCCAGGGTGTTGG | This study |
| | pmrA_rev | CGTGCGATAGCGTGACAACC | This study |
| | tus_fwd | AACAGTCCGAAAATCGCAGC | This study |

| | | |
|---|---|---|
| tus_rev | GCAATCAGTGGTGTAGGGCA | This study |
